# Normal Aging Limits Cortical Network Reorganization and Behavioral Recovery after Experimental Stroke

**DOI:** 10.64898/2026.04.24.720447

**Authors:** Asher J. Albertson, Ryan M. Bowen, Kyrillos Ayoub, Rose A. León-Alvarado, Brendon Wang, Roman Pati, Adam Q. Bauer, Jin-Moo Lee

**Author notes:** These authors contributed equally to this work (Asher J. Albertson, Ryan M. Bowen). **Contact Information:** Jin-Moo Lee: Washington University School of Medicine, 660 S Euclid Avenue, Campus Box 8111, St. Louis, MO 63110.

## Abstract

Stroke is the leading cause of chronic disability in the United States, and advancing age is associated with worse recovery. Despite this, relatively little is known about how aging influences the repair and reorganization of neural circuits and large-scale cortical networks after stroke. To address this question, we compared cortical network dynamics and behavioral recovery after focal photothrombotic stroke in forepaw somatosensory cortex in young (3-month-old) and aged (18-month-old) Thy1-GCaMP6f mice. Both young and aged mice developed significant somatomotor deficits after stroke; however, only young mice exhibited substantial behavioral recovery despite similar infarct volumes across groups. Two age-dependent effects on cortical network function emerged. First, somatosensory-evoked activity and somatosensory functional connectivity were disrupted in both cohorts early after stroke, but their trajectories diverged over time. Forepaw-evoked GCaMP responses in the affected cortex were similarly reduced in both groups early after stroke; yet by 7 weeks, responses recovered in young mice but remained persistently depressed in aged animals. Likewise, bihemispheric somatosensory functional connectivity was initially disrupted in both groups but improved between 1 and 7 weeks only in young mice. Second, global temporal measures of network function evolved differently after stroke. At baseline, stimulus-locked fidelity and interhemispheric coherence were higher in young than aged mice, but after stroke, these measures declined in young animals to levels comparable to aged mice and did not recover by 7 weeks. Stroke also altered large-scale cortical entrainment dynamics, and reductions in cortical entrainment area between baseline and 1-week post-stroke predicted long-term behavioral recovery across animals. Together, these findings indicate that impaired behavioral recovery in aged mice reflects a failure of damaged somatosensory networks to reorganize, whereas recovery in young mice occurs despite persistent degradation of global network fidelity and coherence. These results highlight age-dependent mechanisms of circuit repair after stroke and suggest a potential network-level basis for chronic deficits in stroke survivors.

**Graphical Abstract:** 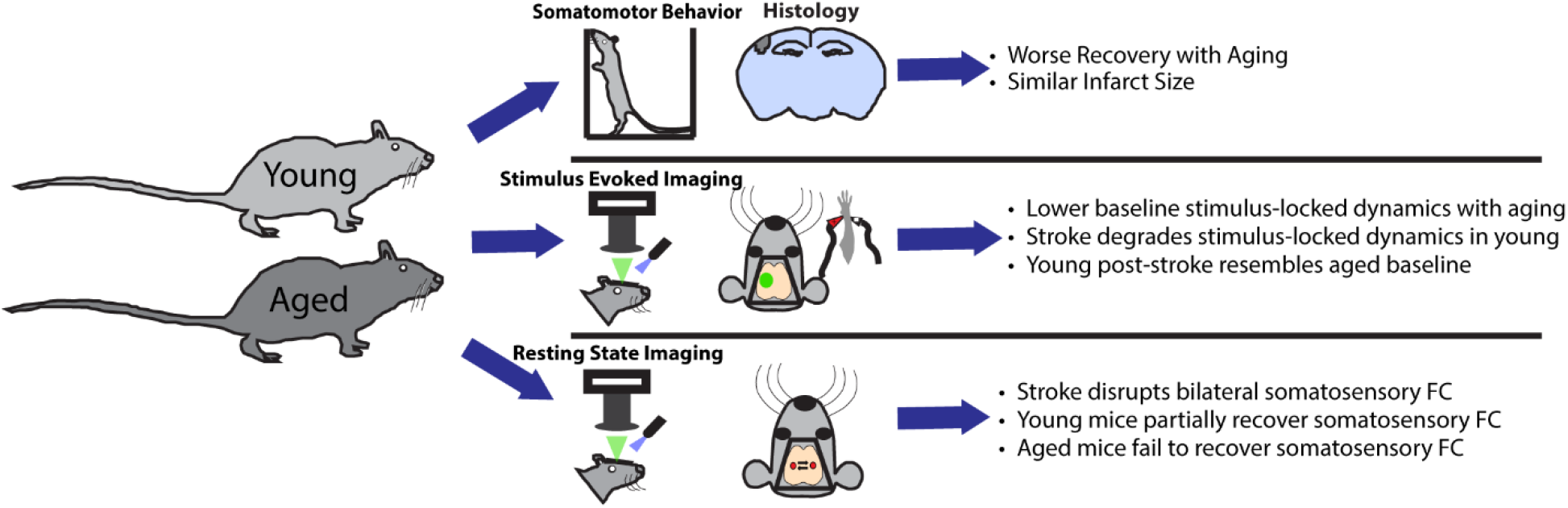

## Introduction

Stroke occurs disproportionately in older individuals ^1^ with age being the greatest non-modifiable risk factor for stroke ^2-4^. This correlates with age-dependent increases in the incidence of stroke risk factors including hypertension^5^ and dyslipidemia^6^. Aging is also associated with dramatically worse outcomes after stroke including increased mortality^7^ and significantly diminished functional recovery^7,8^. The risk of unfavorable neurologic outcomes increases linearly with age^9^. Older age is a negative predictor of functional independence one year after stroke^10^, and older patients are less likely to benefit from specialized stroke rehabilitation units^11^. Importantly, the association between age and worse stroke outcomes occurs independent of stroke severity^12^. Finally, in long term studies of stroke patients, the young cohorts demonstrated continued recovery at 24 months post-stroke and stability at 30 months, while the aged cohorts had more limited recovery and exhibited functional decline as early as 18 months post-stroke^13,14^. While many age-related factors likely contribute to worse recovery after stroke, a diminished capacity of the aged brain for adaptive reorganization of relevant circuits and networks after injury may drive limited functional recovery.

Normal aging is associated with a wide range of changes to the substrate and function of the brain. Brain weight declines ^15^, white matter tract myelin decreases ^16^, and the cerebral cortex thins^17^. At a neuronal level, aging is associated with alterations in dendritic spines^18^ and arbors^19^ as well as changes in axonal bouton dynamics^20^. Both short term plasticity^21^ and long-term potentiation^22^ change during normal aging. The changes in the substrate and cellular behavior of the brain are associated with changes in network level activity. Whole cortex spectral power declines with age^23,24^ particularly in lower frequencies and the degree to which this activity declines correlates with cognitive performance. Cortical activation patterns become more bilateral and less hemispherically specialized with aging^25,26^. Finally, normal aging is associated with a variety of alterations to network dynamics^27,28^ and connectivity^24,29^. These age-related changes together may limit the capacity of the brain to recover after a stroke.

Regional and brain-wide reorganization is a critical component of stroke recovery^30-33^. Recovery from small cortical strokes is associated with remapping of the function of the damaged region to adjacent cortex^34-36^ as well as more distant regions^37,38^. Disruption of cortical remapping limits stroke recovery^39^. At a broader level, stroke is associated with disruption of brain-wide functional networks^40-42^. These disruptions may predict deficits^43,44^ and in particular may predict impairment of less localizable deficits including visual and verbal memory^43^. Restoration of network connectivity is associated with better functional recovery after stroke ^45,46^. Given the alterations in neuronal operation and network function with aging^47,48^, we have hypothesized that aging would be associated with a more limited capacity for network re-organization after stroke and that this may correlate with diminished behavioral recovery.

To address this question, we examined behavioral recovery and cortical network activity in young and aged mice before and during recovery from focal stroke. We focused on forepaw somatosensory cortex, associated somatomotor networks, and spontaneous somatosensory behavior. We measured somatosensory-evoked activity, somatomotor functional connectivity, and broader temporal features of network dynamics, including stimulus-locked fidelity and interhemispheric coherence during peripheral stimulation, and related these measures to behavioral recovery.

## Methods

### Experimental Design

Mice were separated into equal cohorts of aged (n=20) and young (n=20) animals. Young mice were between 3 and 5 months old and aged mice were between 18 and 20 months old during the execution of the experiment. Cohorts were split evenly between males and females. 1 week prior to the experiment, mice were placed in cages with toys and exercise wheels to promote recovery. Each experimental week, behavior analysis was performed at the beginning of the week, and imaging was performed at the end of the week. All mice underwent behavior at baseline, and 1, 3, and 7 weeks after photothrombosis. All mice underwent imaging at baseline and 1 and 7 weeks after photothrombosis (Figure 1). At the conclusion of the experiment, brains were collected for histology from all surviving mice.

**Figure 1.**
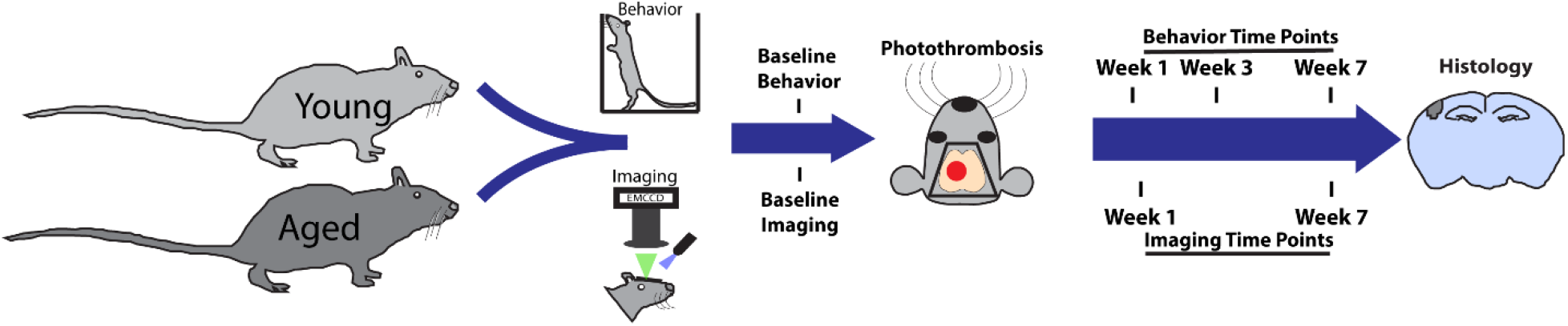
Overview of experimental Design.

### Mice and Procedures

#### Ethical approval

All procedures delineated below were conducted in compliance with the policies of the Washington University Animal Studies Committee and in accordance with the guidelines set forth by the American Association for Accreditation of Laboratory Animal Care.

#### Mouse model

Mice were housed in enriched environments (as previously described^45^) to enhance the dynamic range of post-stroke recovery and were provided food and water ad libitum under a 12-hour light/dark cycle. Experiments used 12-week-old (young) and 18-month-old (aged) hemizygous Thy1-GCaMP6f mice on a C57BL/6J background^49^. Genotype was verified via PCR using the following primers: forward 5’-CATCAGTGCAGCAGAGCTTC-3’ and reverse 5’-CAGCGTATCCACATAGCGTA-3’.

#### Cranial windows for optical imaging

Cranial windows were implanted over the dorsal skull of each mouse following established protocols^50^. Mice were anesthetized with isoflurane (3–5% for induction, 1–2% for maintenance) and secured in a stereotaxic apparatus. Body temperature was regulated using a feedback-controlled heating pad. After shaving and sterilizing the scalp, a midline incision was made, and the skin was retracted. A custom Plexiglas window was affixed to the skull using Metabond dental cement (C&B Metabond, Parkell Inc.), fully covering the surgical area. Mice were allowed a 5-day recovery period prior to any behavioral or imaging assessments.

#### Photothrombosis

Photothrombotic stroke was induced as previously described^39^. Mice received an intraperitoneal injection of Rose Bengal. A 1-mm-diameter region centered in right forepaw somatosensory cortex (left hemisphere; −2.2 mm from bregma, 0.5 mm from lambda) was then illuminated with a 523-nm diode-pumped solid-state laser for 10 minutes. Behavioral and imaging assessments began one week after stroke.

#### Behavioral assays

Following stroke to primary somatosensory forepaw cortex (S1_FP_), mice typically show asymmetric forepaw use favoring the unaffected limb during the cylinder rearing task, which generally becomes more symmetric as recovery progresses^39,51^. Mice were recorded for 10 minutes while engaging in spontaneous rearing behavior. The number of video frames in which each forepaw contacted the cylinder wall was counted to calculate forepaw asymmetry using the formula: Asymmetry = (Left − Right)/(Left + Right + Both), as previously described (Figure 1C)^52^. Behavioral recovery was quantified as the change in asymmetry from Week 1 to Week 7, representing the shift from maximum to chronic deficit. Recovery was defined as any improvement in asymmetry (increased use of the affected paw), while deficit was defined as a worsening of asymmetry (increased reliance on the unaffected paw). All asymmetry values were corrected for baseline asymmetry.

#### Infarct volume quantification

Infarct volumes were measured as previously described^53^. Mice were deeply anesthetized with pentobarbital and transcardially perfused with heparinized phosphate-buffered saline. Brains were extracted, frozen, and sectioned coronally at 40-micron thickness using a sliding microtome. Consecutive sections containing infarcted tissue were mounted and stained with cresyl violet. Brightfield images were captured using a Keyence BZ-X800 microscope. Infarcted areas were outlined and quantified by blinded experimenters using ImageJ. The total infarct volume was calculated by summing the infarct areas across all sections and multiplying by slice thickness. Mice whose strokes were not visible in the final analysis were excluded.

### Wide-field Optical Imaging Recordings

#### Wide-field optical imaging

Wide-field optical imaging (WFOI) was used to measure cortical calcium and hemoglobin signals, as previously described^50^. The dorsal neocortex was sequentially illuminated with three LEDs centered at 470 nm, 525 nm, and 625 nm. GCaMP was excited using the 470 nm LED, while the remaining wavelengths were used for multispectral hemodynamic imaging. Multiwavelength dichroic beam combiners (LCS-BC25-0480, LCS-BC25-0560, LCS-BC25-0585, LCS-BC25-0605; Mightex Systems) were utilized to collimate and merge all LED sources for skull illumination. A cooled sCMOS camera (Zyla 5.5-USB3, Andor) equipped with a 75-mm f/1.8 lens captured images at 80 Hz (25 Hz per channel).

#### Optical imaging recordings

Wide-field optical imaging (WFOI) was conducted as previously^53^. Resting-state imaging to assess functional connectivity was performed for 10 minutes in awake mice. Mice were then anesthetized with an intraperitoneal injection of ketamine-xylazine (100 mg/kg ketamine, 10 mg/kg xylazine) to perform evoked-response imaging during electrical stimulation of the forepaws. Body temperature was maintained as previously noted. Forepaw evoked response imaging was performed on each forepaw for 5 minutes using computer-controlled 1 mA pulses transmitted via microvascular clips (Roboz) placed on either side of the paw. Stimulation followed a block design: 5 seconds rest, 5 seconds of 3 Hz stimulation (1 mA, 0.3 ms pulses), followed by 10 seconds rest, repeated for 15 blocks over the 5-minute session. Imaging sessions were conducted before stroke and at 1 and 7 weeks after stroke.

### Optical Imaging Processing and Analyses

#### Optical imaging signal processing

Brain masks were manually traced for each mouse, and all subsequent analyses were restricted to pixels identified as brain tissue. Brain masks and optical imaging sequences were aligned to Paxinos atlas space using affine transformations based on cranial landmarks^54^. To eliminate background sensor counts, five seconds of dark frames were collected at the beginning of each imaging session and subtracted from all subsequent frames. Spatial and temporal detrending were applied to the time series of all brain pixels as previously described^55^. Oximetric changes were calculated^56^, and hemoglobin-related signal contamination was removed from the GCaMP data using established methods^57^. Global signal regression was applied to all brain pixels to reveal the spatial organization of functional networks. Imaging sessions exhibiting motion artifacts, identified by thresholding fluctuations in light intensity^50^, were excluded from further analysis.

#### Imaging derived infarct areas

Imaging-derived infarct areas were quantified using a previously described approach by Bowen et al.^52^. Resting-state recordings were filtered (1–4 Hz) to generate homotopy maps defined as Pearson z-correlations between mirror-image pixels across the midline. For each mouse, week-1 post-stroke homotopy maps were divided by baseline maps to obtain ratio homotopy maps. Pixels with ratio z ≤ 0.2 were classified as infarcted, and the largest contiguous region below this threshold was defined as the infarct ROI. Infarct area (mm^2^) was computed from the ROI dimensions and used for group comparisons (Figure 2E,F). Homotopy-determined infarct volumes correlate robustly with histological infarct volumes^52^.

**Figure 2.**
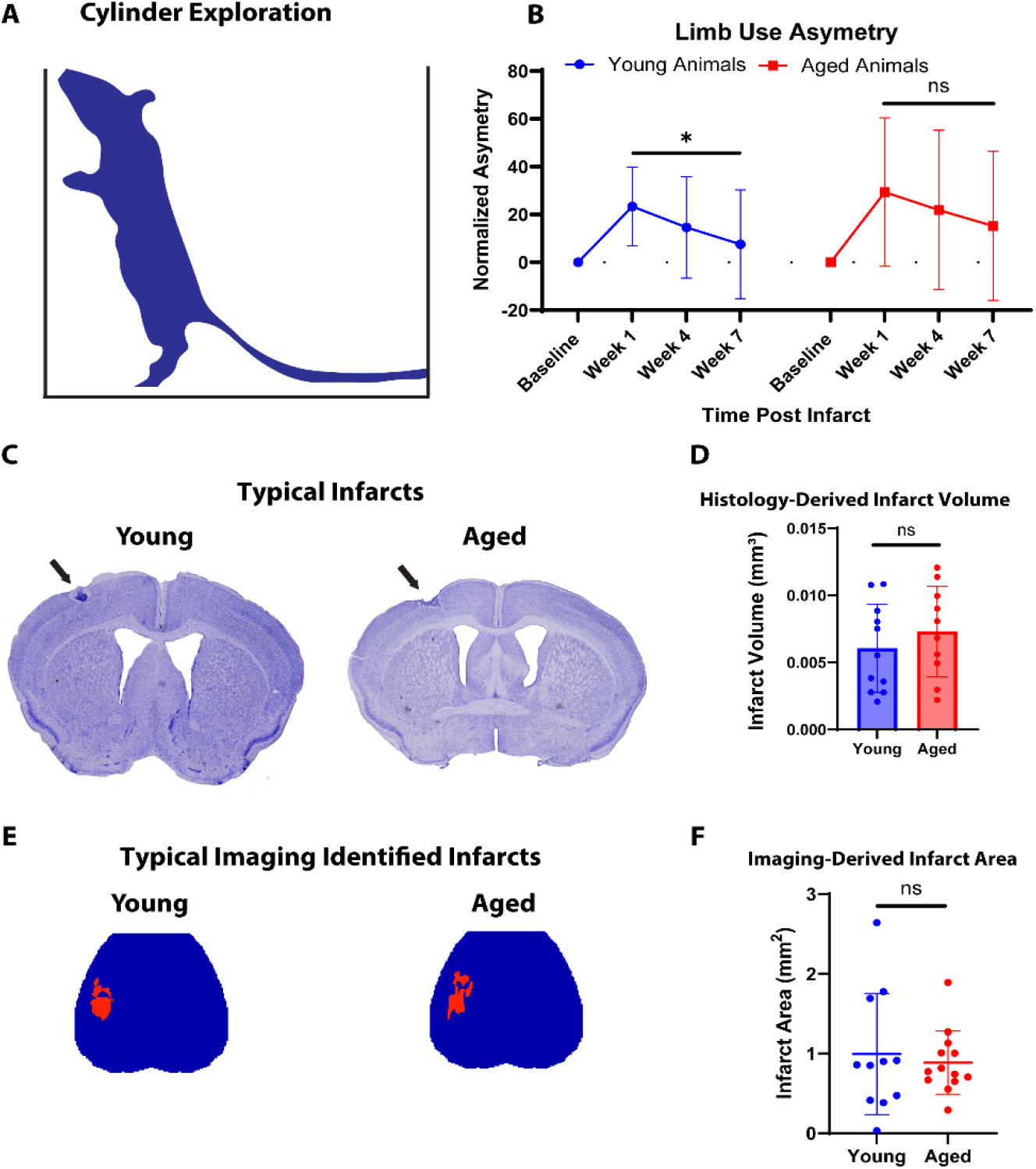
Behavioral Recovery and Infarct Volume. (A,B) Limb-use asymmetry during spontaneous cylinder exploration was normalized to each animal’s baseline value and measured at baseline and 1, 4, and 7 weeks after photothrombotic stroke targeting the forepaw somatosensory cortex. Both young and aged mice developed significant forelimb-use asymmetry after stroke, with no difference in the magnitude of deficit at week 1 between groups. Mixed effects analysis (factors: age and time) followed by Tukey post-hoc comparisons demonstrated significant improvement in asymmetry between week 1 and week 7 in young mice, whereas aged mice did not show a statistically significant improvement over the same interval. (C-D) Representative cresyl violet-stained sections (C) and quantification of infarct volume at 8 weeks revealed no significant difference in infarct volume between young and aged cohorts (D; unpaired t-test; young: n=11, aged: n=10). (E-F) Maps quantifying infarct area using a previously validated imaging-based approach at the week-1 time point likewise showed no significant difference in infarct area between young and aged mice (unpaired t-test; young: n=11, aged: n=13). Data are shown as mean ± SD.

#### Evoked response analysis

Regions of interest (ROIs) corresponding to the forepaw somatosensory cortex were defined by thresholding both individual and group-averaged forepaw evoked responses. For each paw and timepoint, GCaMP ROIs were identified by selecting pixels exceeding 50% of the maximum intensity in peak-averaged response maps, averaged across stimulation blocks and peaks. The evoked response maps presented show group-, block-, and peak-averaged fluorescence signals following each electrical pulse (0.3 ms pulse duration delivered at 3 Hz for 5 seconds). Mice whose peak GCaMP response amplitude dropped below 10% of their individual baseline maximum were excluded from analysis.

#### Power and fidelity analyses

Fast Fourier transforms (FFT) were performed using MATLAB’s native fft() function, following previously established methods^52^, on block-averaged stimulation data from affected forepaw stimulation sessions to obtain magnitude maps at frequencies from 0 to 10 Hz. Fidelity maps were generated by calculating the magnitude of the FFT of each pixel at 3 Hz (the stimulation frequency) ±0.05 Hz. For each mouse, a pixel-wise log_10_ transformation was applied to all power values. Median logarithmically transformed fidelity values were taken within each hemisphere of each mouse for hemispheric fidelity quantifications displayed on plots. Ipsilesional forepaw somatosensory fidelity values were obtained by averaging logarithmically transformed power within individually defined ROIs for each mouse.

#### Phase analysis

Fast Fourier transforms, described above, were used to create individual phase maps during 3-Hz stimulation of the affected forepaw. These maps show the phase position of each pixel in the brain space, from phase 0 to 2π during stimulation. Phase was corrected by multiplying each phase quantity by e^-2πi*f*t0^, where f is the frequency of stimulation (3 Hz) and t0 is the start time of the block-averaged stimulation. Noise from these individual phase maps was removed by finding all pixels whose FFT magnitude was not at least 30% of the maximum FFT magnitude on the power maps described above, and filling the phase of those pixels with the phase of the surrounding pixels whose FFT magnitude did reach the 30% threshold. Phases were then unwrapped using a 2D phase unwrapping algorithm available on MATLAB Central File Exchange^58^. Group-averaged phase maps were created by averaging all non-NAN phase values for a given timepoint and group.

#### Planarity analysis

Local planarity maps were created by determining the R^2^ value of a planar wave model in predicting 3-Hz phase angles, described above, in a 21-pixel x 21-pixel window during block-averaged peripheral stimulation. The planar wave model was fit to the equation z = ax + by + c, where a, b, and c are empirically derived constants, and x and y are the spatial dimensions of the phase maps in the window being fit. This window was then slid across the entire brain space to obtain R^2^ values for the central pixel in each window in the cortex, which is the reason for the inability to assess planarity at the edges of the maps. This model was optimized for window size to ensure that plane fitting was temporally resolved, yet robust and not noisy. Pixel-wise R^2^ values were plotted on maps for each individual animal to quantify the degree of variance that could be explained by a planar wave model in the given window of cortex. All pixels with R^2^ values above 0.8 on individual planarity (R^2^) maps were considered to be entrained to peripheral stimulation. Group-averaged planarity maps were obtained by finding the mean R^2^ value of each analogous pixel across a group and timepoint, and group-averaged entrained regions are outlined in black on group-averaged planarity maps.

#### Magnitude-squared coherence analysis

Evoked response coherence maps were generated for each mouse by computing magnitude-squared coherence (using MATLAB’s native mscohere() function) between block-averaged activity in individually defined ipsilesional S1FP regions of interest (see Evoked response analysis) and the activity of each pixel across the brain during stimulation-on periods of right (affected) paw stimulation at 3 Hz. Individual coherence values were calculated by measuring the magnitude-squared coherence at 3 Hz between each mouse’s average GCaMP signal in the affected forepaw ROI (created during right paw stimulation) and the unaffected forepaw ROI (created during left paw stimulation) during stimulation-on periods of right paw stimulation.

#### Functional connectivity analysis

Resting-state functional connectivity (RSFC) analysis was conducted as previously described^50^. Pre-processed resting-state data were bandpass filtered to isolate specific frequency ranges: infraslow (0.01-0.1 Hz), delta (1–4 Hz), and theta (4–7 Hz). For each mouse, the GCaMP fluorescence signal within a defined region of interest (ROI), or seed, was averaged and correlated with the GCaMP signal from every other brain pixel. Pearson z-transformed correlation maps were generated for each time point to visualize longitudinal changes in functional networks. Seed-based connectivity measures, including node degree, were derived using evoked response ROIs from affected and unaffected forepaw stimulation at corresponding time points, and using Paxinos atlas-based ROIs for forepaw primary somatosensory and primary motor regions. Node degree was defined as the number of brain pixels exhibiting a Pearson z-correlation of ≥ 0.4 with the ROI-averaged GCaMP signal.

### Statistical Analyses

#### Sample size calculation

Mice that died at any point of the experiment were excluded from further analyses. Mice that exhibited motion artifacts or poor connection of forepaw microvascular clips at any timepoint were also excluded from resting state or evoked response analyses, respectively, at that timepoint. Additionally, mice who did not appear to have a stroke based on imaging criteria or post-hoc histological analysis (figure 2) were excluded. The final sample sizes for statistical analyses of cylinder rearing asymmetry were as follows: young group (Week 0: n = 15, Week 1: n = 14, Week 4: n = 14, Week 7: n = 14), aged group (Week 0: n = 15, Week 1: n = 15, Week 4: n = 15, Week 7: n = 15). The final sample sizes for statistical analyses on evoked response optical imaging measures were as follows: young group (Week 0: n = 14, Week 1: n = 18, Week 7: n = 18), aged group (Week 0: n = 16, Week 1: n = 17, Week 7: n = 13). The final sample sizes for statistical analyses on resting state functional connectivity optical imaging measures were as follows: young group (Week 0: n = 10, Week 1: n = 14, Week 7: n = 15), aged group (Week 0: n = 15, Week 1: n = 14, Week 7: n = 15).

#### Statistical analysis

All statistical analyses were conducted using GraphPad Prism version 11.0. All datasets analyzed passed the Shapiro-Wilk test for normality. Descriptive statistics shown on figures are all mean values with error bars showing standard deviation. Longitudinal changes in optical imaging and behavior were analyzed using a two-way mixed-effect model with factors of age (young vs aged) and time. Post hoc Tukey’s tests were applied for multiple comparisons, comparing between age groups at each time point and across time points within age groups. For behavioral limb-use asymmetry measurements, normalized asymmetry values (asymmetry minus baseline asymmetry, were used to correct for pre-existing preference) were analyzed longitudinally across baseline and post-stroke time-points. In addition, planned within-group comparisons between week 1 and week 7 were performed to assess recovery within groups. Infarct volumes were compared between young and aged groups using an unpaired two-tailed Student’s t-test. A p-value of < 0.05 was considered statistically significant for all tests.

## Results

### Aged Animals Have Worsened Recovery Profiles Despite Similar Infarct Sizes

There was no statistically significant difference in the magnitude of the behavioral deficit between young and aged mice at week 1. Consistent with prior findings^59^, young mice demonstrated a significant improvement in limb-use asymmetry between weeks 1 and 7 after stroke (p = 0.0462; Figure 2B). In contrast, aged mice did not demonstrate a statistically significant improvement in limb-use asymmetry between weeks 1 and 7 (Figure 2B). These age-dependent differences in recovery occurred despite infarct size being similar across ages. Infarct volumes assessed histologically at week 8 showed no statistically significant differences between young and aged cohorts (Figure 2C,D). Infarct volumes measured many weeks after ischemia are often confounded by remodeling and atrophy^60,61^, and therefore, we also performed a previously validated^52^ imaging-derived analysis of infarct area at the week 1 time point. This analysis, based on loss of interhemispheric homotopy during resting state imaging (see methods), also showed no difference in stroke size between the aged and young cohorts (Figure 2E,F).

### Forepaw-evoked Cortical Responses Differ Between Young and Aged Mice and Show Distinct Post-Stroke Time Courses

Our group and others have previously shown that photothrombotic strokes to the forepaw representation of the somatosensory cortex in young mice results in an initial deficit followed by restoration of forepaw evoked responses and remapping onto surrounding regions in concert with behavioral recovery^37,39,52^. We used stimulus-evoked (Figure 3A), block-averaged GCaMP fluorescence maps to examine evoked response amplitude during forepaw stimulation in young and aged animals at baseline, and at 1 and 7 weeks after photothrombotic stroke. Evoked response amplitude was quantified as summed fluorescence within the weekly activation ROI (see methods, Figure 3C) and analyzed using a mixed-effects model with Tukey’s post-hoc comparisons. In this cohort, aged animals showed significantly lower GCaMP responses to stimulation of the affected paw at baseline (p = 0.0003; Figure 3B,C). In keeping with prior work, forepaw evoked responses in young animals were significantly reduced at week 1 relative to baseline (week 0 vs week 1: p = 0.0111; Figure 3B,C). Notably, the difference between young and aged animals was no longer statistically significant at week 1 (Figure 3B,C), but re-emerged by week 7, when aged animals again exhibited significantly lower evoked response amplitudes than young (p = 0.0108; Figure 3B,C).

**Figure 3.**
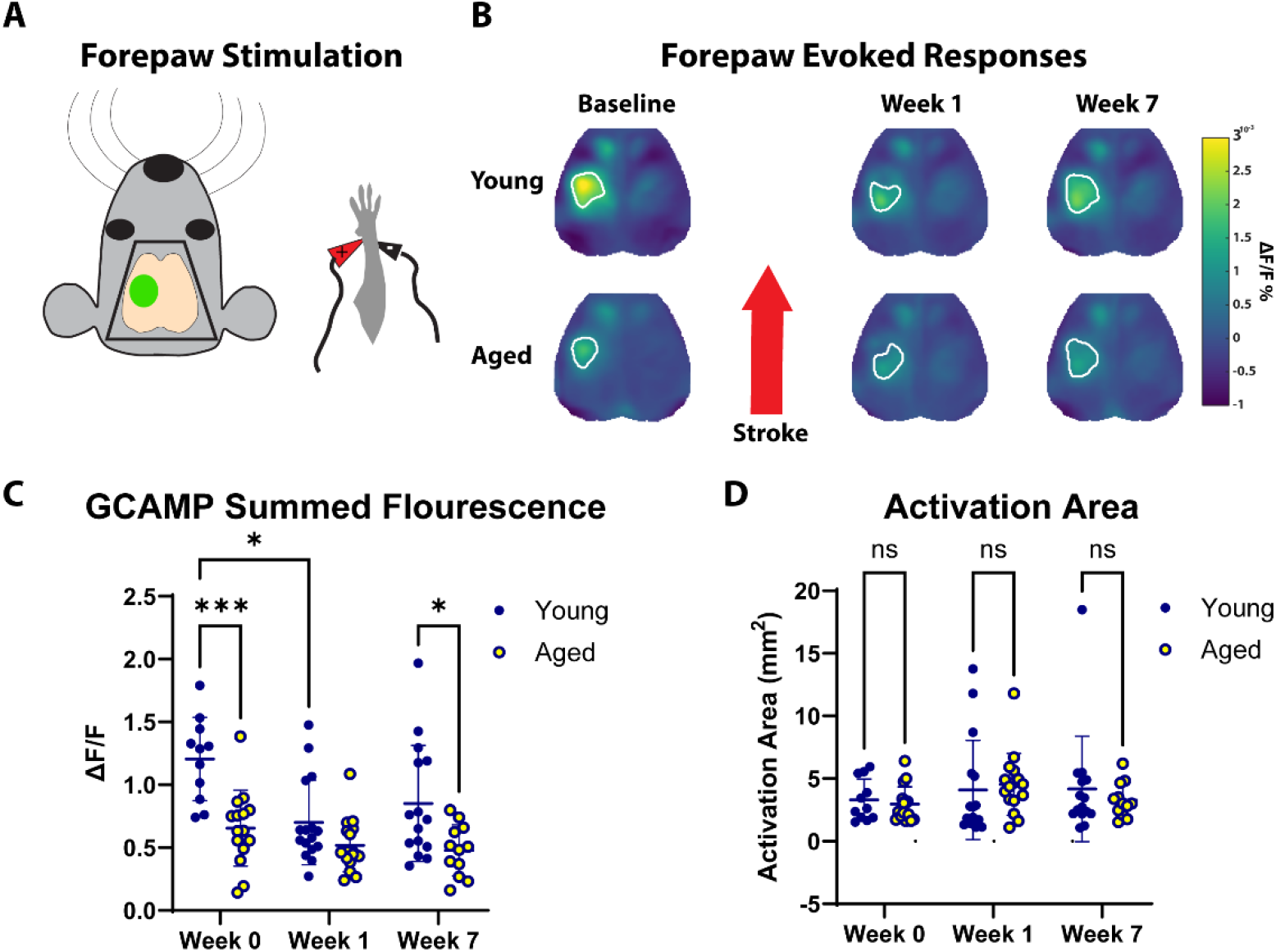
Age-Dependent Difference in Forepaw Evoked Responses After Stroke. (A) Forepaw stimulation was used to evoke somatosensory responses in the infarcted forepaw representation of somatosensory cortex. (B) Representative block-averaged forepaw-evoked GCaMP response maps are shown for young (top row) and aged (bottom row) mice at baseline and at 1 and 7 weeks after photothrombotic stroke. (C) Peak evoked response amplitude (max ΔF/F) and (D) evoked activation area (GCaMP ROI area, mm^2^) are quantified. Mixed-effects analysis (factors: age and time) followed by Tukey post-hoc comparisons demonstrated significantly lower evoked response amplitude in aged compared with young mice at baseline and at week 7, while the between-group difference was not statistically significant at week 1. Evoked activation area did not differ between young and aged mice at baseline or at post-stroke timepoints (ns). Data are shown as mean ± SD. Young cohort sample sizes – Week 0: n = 14, Week 1: n = 18, Week 7: n = 18. Aged cohort sample sizes – Week 0: n = 16, Week 1: n = 17, Week 7: n = 13.

Within-group comparisons did not demonstrate a statistically significant change from week 1 to week 7 in either young or aged animals. In contrast, aged animals remained relatively low across timepoints (Figure 3B,C). Similar to our prior work we did not observe a difference in total GCaMP activation area at baseline or at any timepoint^24^. In summary these results demonstrate age-dependent attenuation of forepaw-evoked response amplitude, with stroke transiently reducing young animals to values not significantly different from aged at week 1, and a significant separation between young and aged again evident by week 7.

### Focal Stroke Disrupts Temporal Dynamics of Neuronal Circuits and Networks in Young and Aged Mice Differentially

Recent work in young mice has shown that stroke disrupts the temporal processing of sensory input, reflected by reduced fidelity of time-locked neuronal responses and decreased bilateral coherence during peripheral stimulation. The magnitude of this disruption correlates with long-term functional outcomes^52^. We therefore examined the fidelity and bilateral coherence of somatosensory cortical responses to peripheral stimulation in aged and young mice following stroke.

To quantify circuit fidelity, we measured stimulus-locked power of forepaw-evoked responses during wide-field calcium imaging. Specifically, 3-Hz stimulus-locked GCaMP fluorescence power across the dorsal neocortex during rhythmic forepaw stimulation was used as a measure of somatosensory circuit fidelity. At baseline, young mice exhibited significantly greater stimulus-locked 3-Hz power (fidelity) than aged mice during rhythmic forepaw stimulation in both the ipsilesional (p = 0.0002; Figure 4B,C) and contralesional hemisphere (p = 0.0007; Figure 4B,D), consistent with stronger entrainment of cortical activity. Aged mice exhibited low stimulus-locked power in both hemispheres, and these values remained low throughout the post-stroke period (Figure 4B-D). Following stroke, stimulus-locked power decreased in young mice both ipsilesionally (week 0 vs week 1: p = 0.0008, week 0 vs week 7: p = 0.0004; Figure 4B,C) and contralesionally (week 0 vs week 1: p = 0.0024, week 0 vs week 7: p = 0.0049; Figure 4B,D), while aged mice exhibited consistently low responses across time points (Figure 4B-D). In ipsilesional forepaw somatosensory cortex, young and aged mice demonstrated comparable levels of fidelity at all time points. Fidelity measures in ipsilesional forepaw somatosensory cortex decreased significantly in both groups at 1 and 7 weeks after stroke (young: week 0 vs week 1: p = 0.0042, week 0 vs week 7: p = 0.0011; aged: week 0 vs week 1: p = 0.0081, week 0 vs week 7: p = 0.0015; Figure 4B,E).

**Figure 4.**
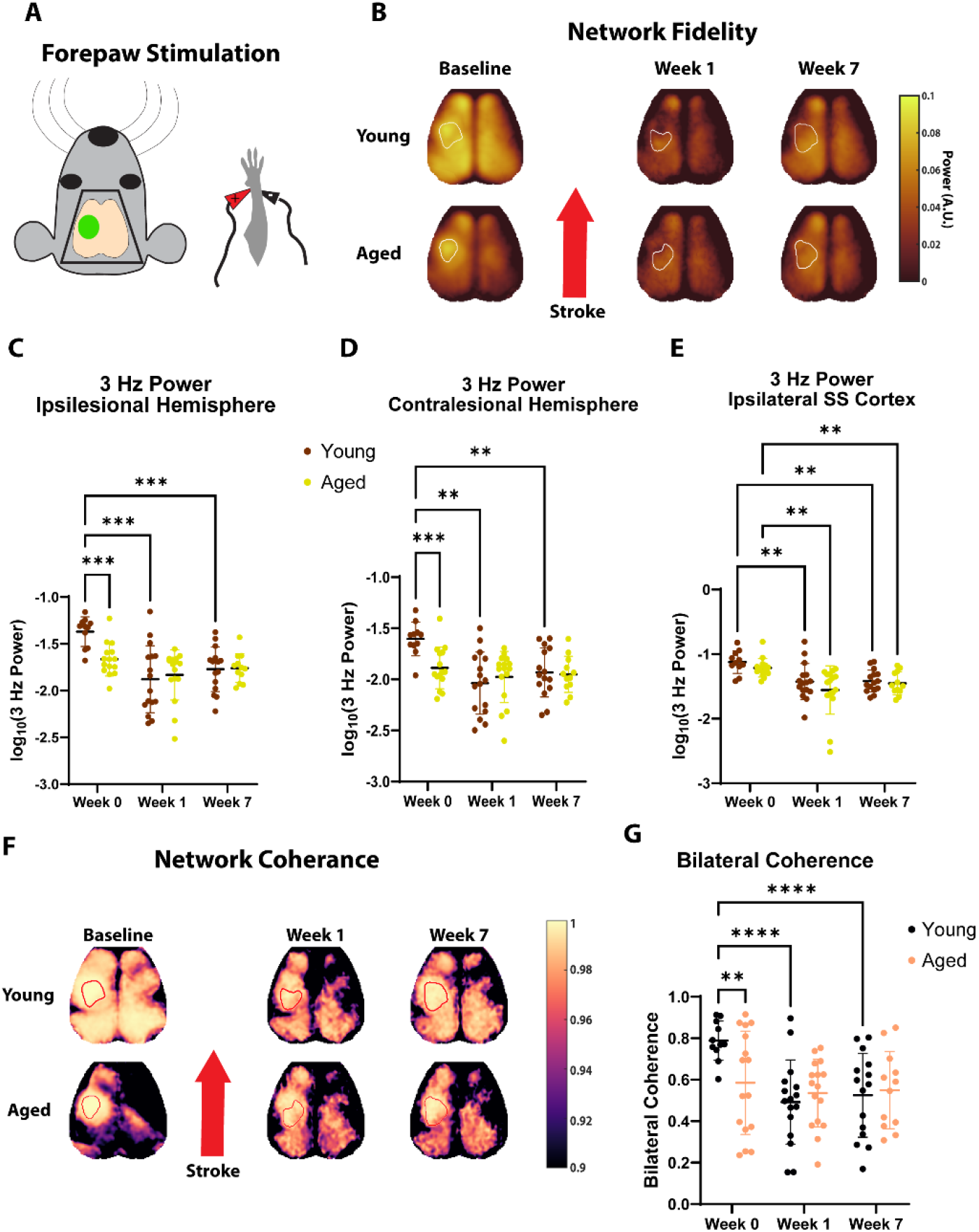
Focal Stroke Reduces Stimulus-Locked (3-Hz) Cortical Power and Bilateral Coherence During Forepaw Stimulation. (A) Neuronal coherence was quantified as stimulus-locked 3-Hz power during stimulation of the affected forepaw. (B) Group-averaged 3-Hz power maps across the cortex during stimulation are shown at baseline and at 1 and 7 weeks after photothrombosis in young and aged mice. (C-E) Quantification of average 3-Hz power within the ipsilesional hemisphere (C), the contralesional hemisphere (D), and the ipsilesional somatosensory cortex (E) is shown. (F-G) Bilateral coherence between forepaw ipsilesional and contralesional somatosensory cortices was quantified as 3-Hz magnitude-squared coherence during stimulation of the affected forepaw. (F) Group-averaged coherence maps across the cortex during stimulation are shown at baseline and at 1 and 7 weeks after photothrombosis in young and aged mice. (G) Quantification of bilateral coherence between somatosensory forepaw regions is shown. Data are shown as mean ± SD and were analyzed using a mixed-effects model with factors of time and age (p < 0.05). Sample sizes for all analyses were as follows: young cohort-Week 0: n = 14, Week 1: n = 18, Week 7: n = 18. Aged cohort-Week 0: n = 16, Week 1: n = 17, Week 7: n = 13.

We applied a similar approach to examine changes in the temporal dynamics of global brain networks after stroke, using 3-Hz magnitude-squared coherence between ipsilesional and contralesional somatosensory forepaw circuits. Magnitude-squared coherence of global brain networks has been shown to be a sensitive measure of neuronal network function, predicting functional recovery after stroke^52^. At baseline, young mice exhibited significantly greater bilateral coherence than aged mice (p = 0.0020; Figure 4F,G). Following stroke, magnitude-squared coherence significantly decreased in young mice (week 0 vs week 1: p < 0.0001, week 0 vs week 7: p < 0.0001; Figure 4F,G), whereas aged mice showed no significant change across time points. As a result, coherence values in young mice after stroke were similar to those observed in aged mice both before and after stroke (Figure 4G).

We also investigated changes in cortical entrainment, a measure of cortical neuronal propagation dynamics associated with recovery after stroke^62^, by quantifying local phase planarity. This was accomplished by determining the variance explained (R^2^) by fitting a planar traveling-wave model to local windows of phase maps (Figure 5A) generated from forepaw-evoked responses using a fast Fourier transform. Using this approach, we quantified the spatial extent of cortex whose phase dynamics were well described by this planar model, representing cortical regions entrained to peripheral stimulation. Our analysis revealed that the cortical area entrained to stimulation was significantly smaller in aged mice than in young mice at baseline (p = 0.0101; Figure 5B,C). Following stroke, the spatial extent of entrained cortex decreased in young mice (week 0 vs week 1: p = 0.0002, week 0 vs week 7: p < 0.0001; Figure 5B,C) but remained statistically unchanged in aged mice. Finally, linear regression analysis demonstrated that the change in cortical entrainment area from baseline to one week after stroke significantly predicted long-term functional recovery across age groups (R^2^ = 0.1835, p = 0.0467; Figure 5D).

**Figure 5.**
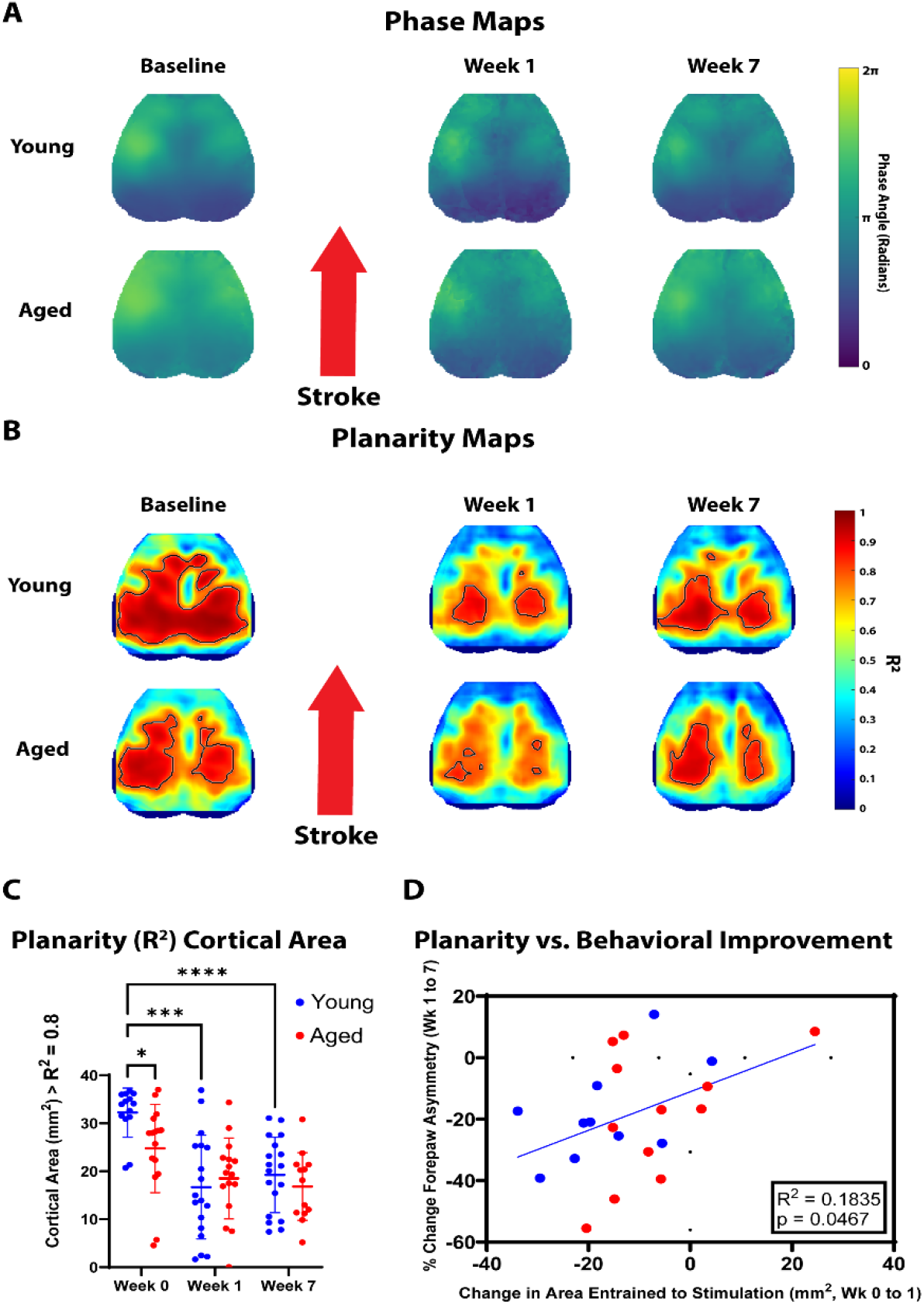
Focal stroke alters cortical entrainment to peripheral stimulation. (A) Phase maps during stimulation across the cortex at baseline and 1 and 7 weeks after stroke in young and aged mice. (B) Local phase planarity maps (R^2^), representing the variance explained by fitting a planar traveling-wave model to local phase windows. (C) Quantification of the cortical area exhibiting strong entrainment. Data are shown as mean ± SD and were analyzed using a mixed-effects model with factors of time and age. At baseline, young mice exhibited greater entrainment area than aged mice (p < 0.05). Following stroke, entrained cortical area decreased in young mice, whereas aged mice showed no significant change across time points (p > 0.05). (D) Linear regression demonstrated that the change in cortical entrainment area from baseline to one week after stroke predicted long-term functional recovery across age groups (R^2^ = 0.18, p < 0.05). Young cohort sample sizes – Week 0: n = 14, Week 1: n = 18, Week 7: n = 18. Aged cohort sample sizes – Week 0: n = 16, Week 1: n = 17, Week 7: n = 13.

### Resting-State Functional Connectivity Reveals Age-Dependent Differences in Network Recovery After Stroke

To investigate longitudinal spatial changes in large-scale neuronal networks, we analyzed region-based resting-state functional connectivity between cortical regions of interest (Figure 6). Homotopic connectivity between bilateral forepaw somatosensory cortices differed in its trajectory between age groups (Figure 6B). Following stroke, connectivity decreased in both young (p < 0.0001; Figure 6A,B) and aged mice (p = 0.0002; Figure 6A,B) at week 1 compared to baseline. However, aged mice showed no significant recovery in connectivity between weeks 1 and 7, whereas young mice demonstrated a significant increase in homotopic connectivity during this period (p < 0.0389; Figure 6A,B). In contrast, ipsilesional connectivity between forepaw somatosensory and motor cortices increased after stroke in both young and aged mice (young: p = 0.0088, aged: p = 0.0017; Figure 6A,C). However, the post-stroke time course differed between groups, with somatomotor connectivity significantly decreasing to more normal levels from week 1 to week 7 in young mice (p = 0.0174; Figure 6A,C) but not in aged mice.

**Figure 6.**
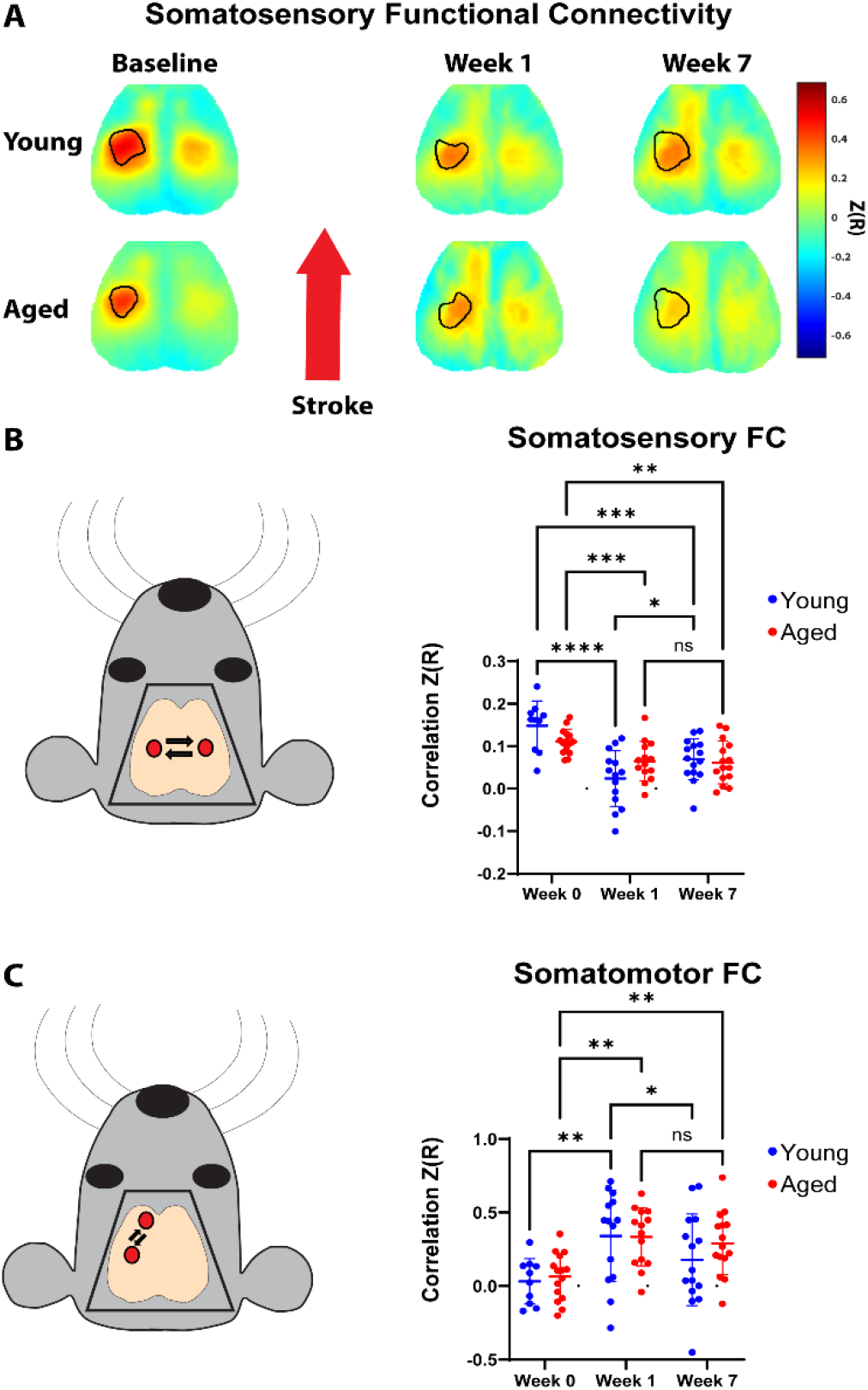
Resting-state functional connectivity reveals age-dependent differences in network recovery after stroke. (A) Functional connectivity maps showing correlation between the functionally identified ipsilesional forepaw somatosensory cortex and the rest of the cortex at baseline and at 1 and 7 weeks after photothrombotic stroke in young and aged mice. (B) Bilateral functional connectivity between homotopic forepaw somatosensory cortices. Young mice exhibited a reduction in homotopic connectivity after stroke followed by partial recovery by week 7, whereas aged mice showed reduced connectivity without recovery. (C) Ipsilesional functional connectivity between forepaw somatosensory and motor cortices. Somatomotor connectivity increased after stroke in both young and aged mice. Data are shown as mean ± SD and were analyzed using a mixed-effects model (p < 0.05). Young cohort sample sizes – Week 0: n = 10, Week 1: n = 14, Week 7: n = 15. Aged cohort sample sizes – Week 0: n = 15, Week 1: n = 14, Week 7: n = 15.

## Discussion

In this study, we used longitudinal behavioral testing and widefield imaging to show that aging is associated with diminished recovery after focal stroke, as well as baseline network dysfunction and persistent impairments in post-stroke circuit and network reorganization. Consistent with prior work, young mice developed a reliable sensorimotor deficit after S1FP photothrombosis with significant improvement at seven weeks post-stroke^39^. Aged mice, despite a similar magnitude of acute impairment, failed to significantly recover during the same interval. This pattern is consistent with human stroke outcomes, in which increasing age is strongly associated with reduced functional recovery^7,8^. Prior preclinical work using transient middle cerebral artery occlusion (tMCAO) (which produces substantially larger, mixed cortical-subcortical infarcts) has similarly shown that aged animals display worsened recovery profiles accompanied by increased gliosis and microglial activation^63,64^. Our work extends this observation to focal somatosensory photothrombotic strokes. In our data, aged mice failed to recover despite exhibiting initial behavioral deficits comparable to those observed in young mice and no difference in infarct volume between cohorts. Histological infarct volume measured 8 weeks after stroke and a previously validated imaging-based metric of lesion area at Week 1 likewise showed no difference between aged and young cohorts. Together, our findings are consistent with prior evidence demonstrating worse recovery profiles with aging. Our data show that this is despite equal infarct sizes, suggesting that diminished recovery in aged mice is driven by a reduced capacity for post-stroke neuronal reorganization rather than differences in lesion size.

Given this, we next examined whether the worsened recovery profiles in aging were associated with diminished circuit reorganization after stroke. Recovery after cortical stroke is dependent on re-establishment of activity within peri-infarct cortex, including restoration of thalamocortical inputs and recruitment of adjacent somatosensory areas^37,59,65,66^. This process depends on local and network level activity^39,67^, expression of plasticity-related proteins^39,68,69^, and is dependent on stroke size^52^. We assessed thalamocortical circuit reorganization in part by quantifying changes in evoked response amplitudes during peripheral forepaw stimulation. Similar to our prior work^70^, the overall area of activation was comparable between young and aged mice; however, aged mice exhibited reduced baseline activation amplitudes before stroke. Following injury, young mice showed diminished activation at one week, and by seven weeks evoked responses were significantly greater than those observed in aged mice, whereas responses in aged mice remained relatively low across time points.

As amplitude-based calcium fluorescence maps lacked information regarding the dynamics of the neuronal circuits and networks we interrogated, we also assessed temporal properties of cortical network function using stimulus-locked fidelity, bilateral coherence, and cortical entrainment. These three measures provide insight into the capacity of the assayed circuits (in this case, thalamocortical circuits and associated corticocortical networks), to stabilize neural activity during peripheral input^71,72^. Higher levels of fidelity, coherence, and entrainment are hypothesized to be mediated by redundant within-network circuitry, which reduces response variability^73,74^. As such, lower baseline fidelity across both hemispheres of aged mice relative to young mice may reflect age-related degradation of corticocortical network dynamics. In contrast, while aged mice showed reduced baseline amplitudes of evoked somatosensory responses, baseline levels of fidelity restricted to the ipsilesional forepaw somatosensory cortex were similar between young and aged mice, suggesting relative preservation of temporal processing in the local circuit with aging. Collectively, these findings suggest that dynamics of longer-range cortical networks may be more vulnerable to age-related degradation than those of local circuits.

This interpretation is strengthened by our observation that age-related baseline differences were not limited to hemispheric fidelity, but were also evident in bilateral coherence and cortical entrainment. These age-related differences at baseline may be associated with degradation of recurrent horizontal cortical projections that relay information throughout the cortex, as has been documented in the auditory cortex of mice^75^. Consistent with this interpretation, our observation that fidelity, coherence, and entrainment levels in young mice at seven weeks post-stroke were strikingly similar to those of aged mice at baseline suggests that both aging and stroke may drive a common degradation of recurrent network connectivity. Taken together, these results raise the possibility that even small somatosensory strokes can induce long-term and widespread disruptions of neuronal networks in young mice, whereas in aged mice, the dynamics of these networks may already operate at a functional floor and therefore show little additional degradation after stroke. Ultimately, these results may reflect a loss of network resilience with aging.

We also demonstrate changes in evoked cortical entrainment that are influenced by both age and stroke. Cortical entrainment strength to both task-based^76^ and automatic^77^ auditory stimuli has been shown to decrease with aging in humans. Furthermore, the strength and phase structure of oscillatory activity in sensory and motor cortex adjacent to stroke has been shown to be correlated with functional recovery^62,78^. The present study adds to these findings in several ways. We reveal disruptions of baseline cortical entrainment dynamics that occur with normal aging, demonstrate a differential impact of stroke on cortical entrainment in young vs aged mice, and show that reductions in cortical entrainment area from baseline to one week after stroke predict long-term functional recovery across age groups. The latter relationship suggests that the capacity of brain networks to sustain spatially organized stimulus-locked activity may represent an important determinant of behavioral recovery after stroke. Furthermore, it is notable how spatially extensive and durable these disruptions in stimulus-locked cortical entrainment are despite the relatively small size of the focal ischemic injury. These findings suggest that both aging and stroke influence the dynamic flexibility of cortical networks in a functionally relevant fashion.

Another possible contributor to degradation of network fidelity, coherence, and entrainment may be age- and stroke-related disruption of inhibitory neuronal networks. Inhibitory neurons are necessary for entraining neuronal synchrony^79,80^, and parvalbumin-positive interneurons in particular modulate gamma oscillations^81^, which are strongly associated with functional recovery after stroke^82^. Stroke significantly disrupts regional inhibitory currents^83^, and modulating GABAergic activity in stroke-affected circuits impacts functional recovery^83,84^, suggesting a key role for inhibitory neurons in post-stroke repair. Furthermore, deregulation of inhibitory circuits in the somatosensory cortex of aged mice is associated with impaired sensory conditioning^85^, while deregulation of these circuits in rat auditory cortex produces distortions in frequency tuning^86^. It is interesting to consider that the observed degeneration of network dynamics at baseline in aged mice and after stroke in young mice may both be partially due to disruptions in inhibitory network activity.

Functional recovery after stroke is associated not only with cortical remapping, but also with restoration of connectivity within and between somatosensory and motor networks. Human studies have shown that recovery is strongly correlated to restoration of widespread functional networks^43,87,88^. In mice, preventing restoration of bilateral somatosensory functional connectivity after stroke significantly diminishes stroke recovery^67^. Furthermore, aging is independently associated with network-specific changes in functional connectivity^24,29^. We examined resting state functional connectivity across interhemispheric somatosensory networks and ipsilesional somatomotor networks. In our model, both young and aged mice exhibited a marked reduction in bilateral S1FP functional connectivity at one-week post-stroke. However, by seven weeks young mice showed partial recovery of interhemispheric functional connectivity, whereas aged mice showed no significant improvement over the same interval. The persistence of reduced connectivity in aged mice, despite similar infarct sizes and comparable early deficits, suggests that aging diminishes the capacity of large-scale somatosensory networks to re-establish coordinated activity after injury. Interestingly, both young and aged mice exhibited increased connectivity between the ipsilesional primary forepaw somatosensory cortex and the ipsilesional primary forepaw motor cortex after stroke. However, only young mice showed a significant decline in somatomotor connectivity toward baseline levels between weeks 1 and 7, whereas aged mice remained relatively elevated across this interval. This abnormally enhanced somatomotor connectivity has been shown to be maladaptive in humans when it persists following stroke^89^, while sustained disruption of functional sensorimotor networks predicts poor recovery in humans^90-92^ and animal models^54^. Together with the impaired recovery of evoked responses, these findings indicate that both local circuit remapping and broader network-level reintegration are disrupted in aging and may contribute to limited capacity for recovery.

Our work identifies two unique age-dependent processes that shape stroke recovery. The first involves local thalamocortical circuits and somatosensory networks: young mice develop robust deficits in both evoked responses and somatosensory functional connectivity after stroke and show partial recovery of each on the same timescale as their behavioral improvement. Aged mice, in contrast, fail to recover either thalamocortical responsiveness or bilateral somatosensory connectivity and show no behavioral improvement, consistent with an age-related loss of the plasticity-dependent mechanisms required for circuit repair. The second process reflects temporal coordination across cortical networks. Here, young mice developed persistent disruptions in stimulus-locked fidelity, interhemispheric coherence, and cortical entrainment of neuronal networks after stroke, reaching levels indistinguishable from those of aged mice at baseline, while aged mice, already operating at low baseline values, showed little change. These findings suggest that aging constrains mechanisms of network reorganization necessary for recovery, whereas stroke itself induces a long-lasting, “age-like” degradation of global temporal network dynamics even in young animals. Together, these data point toward a dual-process model of stroke recovery in which aging limits the capacity for network-level repair while stroke accelerates network degradation typically associated with aging.

Our study has several limitations. First, our imaging approach is restricted to cortical calcium signals; contributions from subcortical structures, including thalamus and basal ganglia, are not directly assessed. Second, our experiments only extended to 8 weeks as many studies have shown recovery in this timespan^39,93^. However, it is possible that additional repair and recovery occurred after this time frame. Third, our strokes were highly focal and targeted specifically to forepaw somatosensory cortex; the impact of aging on network reorganization and temporal dynamics after strokes affecting other cortical or subcortical systems remains unexplored. Future studies will be needed to define the molecular, structural, and immunologic processes that underlie baseline network dysfunction in aged mice, as well as the mechanisms driving the stroke-induced spatiotemporal degradation observed in young mice. In particular, it will be important to determine whether interventions aimed at restoring peri-infarct plasticity or improving global temporal coordination can differentially rescue these two processes and ultimately enhance recovery across the aging spectrum.

